# Malt1-dependent cleavage of Tensin-3 controls B-cell adhesion and lymphomagenesis

**DOI:** 10.1101/2022.09.29.510036

**Authors:** Mélanie Juilland, Nagham Alouche, Ivana Ubezzi, Montserrat Gonzalez, Tabea Erdmann, Georg Lenz, Sanjiv A. Luther, Margot Thome

## Abstract

The protease Malt1 controls the development and function of lymphocytes and promotes lymphomagenesis by cleaving a limited set of cellular substrates, many of which regulate gene transcription. Here, we report the identification of the integrin-binding scaffold protein Tensin-3 as a Malt1 substrate in activated B cells. B cells expressing a non-cleavable form of Tensin-3 (TNS3-nc) showed normal NF-κB and JNK transcriptional responses but increased and prolonged integrin-dependent adhesion upon activation. Moreover, mice expressing a non-cleavable form of Tensin-3 displayed reduced antibody production in response to immunization with a T-cell dependent antigen. We also explored the role of Tensin-3 in diffuse large B cell lymphomas and mantle cell lymphomas characterized by constitutive Malt1 activity, which showed strong constitutive Tensin-3 cleavage and a correlating reduction in total Tensin-3 levels. Silencing of Tensin-3 expression in Malt1-driven lymphoma models did not affect cellular proliferation but enhanced the dissemination of xenografted lymphoma cells. Thus, Malt1-dependent Tensin-3 cleavage limits integrin-dependent B-cell adhesion and promotes humoral immune responses and metastatic spreading of B cell lymphomas in a transcription-independent manner.

## Introduction

Antigen-dependent triggering of the B-cell receptor (BCR) or T-cell receptor (TCR) induces the formation of multimeric signaling complexes that activate lymphocytes by orchestrating changes in gene transcription, cell adhesion and migration. The arginine-specific cysteine protease Malt1 (also known as paracaspase) has recently emerged as a signaling protein that contributes to several aspects of lymphocyte activation and differentiation by the inducible cleavage of a limited set of cellular substrates (Meininger & Krappmann, 2016; Ruland & Hartjes, 2019). Indeed, mice lacking Malt1 are immunodeficient and show striking defects in the antigen receptor-mediated activation of B- and T-cells and the development of innate-like marginal zone (MZ) and B1 B cells (Ruefli-Brasse *et al*, 2003; Ruland *et al*, 2003). Generation of mice expressing a catalytically inactive form of Malt1 revealed that the protease activity of Malt1 accounted to a large part for the capacity to mount proper B- and T-cell dependent immune responses (Bornancin *et al*, 2015; Gewies *et al*, 2014; Jaworski *et al*, 2014; Yu *et al*, 2015). Like Malt1-deficient mice, mice expressing inactive Malt1 showed impaired development of MZ and B1 B cells. Additionally, both Malt1-deficient and Malt1-inactive mice have a defect in the thymic development of regulatory T (Treg) cells (Bornancin *et al.*, 2015; Gewies *et al.*, 2014; Jaworski *et al.*, 2014; Yu *et al.*, 2015). The Treg defect is without consequences in Malt1 ko mice, which cannot mount B- and T-cell responses, but leads to autoimmunity in Malt1-inactive mice, since the remaining scaffold function of Malt1 allows for partial immune responses that are not efficiently counterbalanced by Treg cells (Bornancin *et al.*, 2015; Gewies *et al.*, 2014; Jaworski *et al.*, 2014; Yu *et al.*, 2015). Collectively, these studies have thus revealed an essential role for the protease activity of Malt1 in the development and activation of various lymphocyte subsets, although the contribution of individual Malt1 substrates to the observed phenotypes remain largely unexplored.

Biochemical studies have provided insight into the regulation of Malt1 protease activity and its contribution to lymphocyte activation. Upon antigen receptor engagement, Malt1 is activated as a consequence of the PKCβ− or PKCθ-mediated phosphorylation of the scaffold protein CARMA1 (also known as CARD11) (Matsumoto *et al*, 2005; Sommer *et al*, 2005), triggering a conformational change in CARMA1 leading to the recruitment of Malt1 via the adaptor protein Bcl10 (Gaide *et al*, 2002; Pomerantz *et al*, 2002; Wang *et al*, 2002). The resulting multimeric Carma1-Bcl10-Malt1 (CBM) complex is thought to favor Malt1-dependent recruitment of additional signaling proteins and to unleash the protease activity of Malt1 by means of Malt1’s inducible oligomerization and monoubiquitination (Pelzer *et al*, 2013; Qiao *et al*, 2013; Schairer *et al*, 2020; Schlauderer *et al*, 2018; Sun *et al*, 2004).

Once activated, Malt1 controls various aspects of lymphocyte activation and differentiation, mainly through the regulation of gene transcription (Meininger & Krappmann, 2016; Ruland & Hartjes, 2019). As a scaffold, Malt1 promotes the activation of the transcription factor NF-κB1 by recruiting the ubiquitin ligase TRAF6, which favors the activation of the IκB kinase (IKK) complex and leads to the phosphorylation-dependent degradation of the NF-κB1 inhibitor IκB by the proteasome (Noels *et al*, 2007; Sun *et al.*, 2004). The protease activity of Malt1, on the other hand, strengthens NF-κB1 activation in an IKK-independent manner, through the autoprocessing of Malt1 (Baens *et al*, 2014) and the cleavage of the deubiquitinating enzyme A20, the NF-κB family member RelB and the LUBAC inhibitory component HOIL-1 (Coornaert *et al*, 2008; Douanne *et al*, 2016; Elton *et al*, 2016; Hailfinger *et al*, 2011; Klein *et al*, 2015). Additionally, Malt1-dependent cleavage of the deubiquitinating enzyme CYLD contributes to the activation of transcription factors of the AP-1 family (Staal *et al*, 2011). Finally, Malt1 protease activity also regulates the gene expression and protein stability of the short-lived transcription factor Myc, which is crucial for the growth of B-cell derived mantle cell lymphoma (Dai *et al*, 2017) and the expansion of Treg cells (Rosenbaum *et al*, 2022).

In addition to activating gene transcription, Malt1 controls gene expression by regulating the stability of mRNA transcripts. Indeed, Malt1 cleaves and thereby inactivates several negative regulators of mRNA stability, including the RNAses Regnase-1 (also known as MCPIP-1 or ZC3H12a) (Uehata *et al*, 2013) and N4BP-1 (Yamasoba *et al*, 2019) and the post-transcriptional repressor proteins Roquin-1 and - 2 (Jeltsch *et al*, 2014). A common denominator of these Malt1 substrates seems to be their capacity to degrade short-lived pro-inflammatory transcripts with conserved secondary RNA structure elements targeting them for degradation (Juilland & Thome, 2018); the half-lives of these short-lived transcripts are thus increased because of Malt1 activation.

Collectively, these observations suggest that Malt1-dependent cleavage of substrates plays an important role in the control of gene expression, either by releasing the brakes on gene transcription or by preventing the degradation of specific sets of pro-inflammatory gene transcripts. However, a limited number of Malt1 substrates have been identified that exert transcription-independent functions, which are linked to the regulation of the cytoskeleton and cell adhesion. Cleavage of the Malt1-binding partner Bcl10, for example, controls cellular adhesion of activated T cells by beta1 integrins (Rebeaud *et al*, 2008). Moreover, cleavage of the tumor suppressor LIM domain and actin-binding protein 1 (LIMA1) by the oncogenic API2-Malt1 fusion protein generates a LIM-domain-only (LMO)-like oncogene that supports proliferation and adhesion of B cells (Nie *et al*, 2015). Finally, cleavage of CYLD, in addition to promoting AP-1-dependent transcription, favors the disassembly of microtubules and induces changes in endothelial cell permeability in response to the vasoactive substance thrombin (Klei *et al*, 2016). Thus, Malt1-dependent substrate cleavage controls various transcription-dependent and - independent aspects of the adaptive immune response and other proinflammatory processes. In the present study, we identified and characterized a new Malt1 substrate, Tensin-3.

Tensins form a family of 4 structurally related proteins encoded by separate genes. Tensin-1, −2 and −3 are thought to regulate the adhesion and migration of epithelial cells, by forming a physical link between the actin cytoskeleton and the cytoplasmic portion of certain beta integrin chains (Blangy, 2017; Lo, 2004). Binding of these Tensins to actin is mediated by an actin-binding domain (ABD) present within the N-terminal region (Lo, 2004). This motif is lacking in the shorter protein Tensin-4 (also known as CTEN, for C-terminal Tensin-like protein). The C-terminal region of all 4 Tensin proteins contains a Src homology-2 (SH2) and a phosphotyrosine-binding (PTB) domain that is thought to interact with NPxY motifs in the cytoplasmic tails of certain integrin beta chains (Calderwood *et al*, 2003; Trub *et al*, 1995). The sequence between the ABD and the SH2 domain is unique to each Tensin family member, suggesting additional, exclusive physiological functions. Indeed, Tensin-3-deficient mice show severe growth defects and postnatal lethality, in contrast to mice lacking Tensin-1 or Tensin-2, which grow normally but exhibit impaired kidney functions in later life (Chiang *et al*, 2005; Cho *et al*, 2006; Lo *et al*, 1997). The early lethality of mice lacking Tensin-3 most likely results from defects in the development of the lung and intestinal epithelium (Chiang *et al.*, 2005).

Here, stable isotope labelling by amino acids in cell culture (SILAC), in combination with a quantitative proteomic approach, was deployed to discover novel Malt1 substrates. We thus identified Tensin-3 as a Malt1 substrate that is cleaved upon B-cell activation and in lymphoma cell lines with constitutive Malt1 activity. B cells expressing a non-cleavable Tensin-3 mutant showed an increased integrin-dependent adhesion response. Moreover, mice expressing a non-cleavable form of Tensin-3 had impaired humoral immune responses, but no defects in T-cell development or function. Finally, Tensin-3 silencing increased the dissemination of xenografted diffuse large B-cell lymphomas (DLBCL). Thus, Tensin-3 acts as a Malt1 substrate that controls the function of normal and transformed B cells by regulating their adhesiveness.

## Results

### Malt1 cleaves the Tensin family member, Tensin-3

We and others have previously shown that Malt1 is constitutively active in cell lines derived from the ABC subtype of DLBCL (Ferch *et al*, 2009; Hailfinger *et al*, 2009). We therefore reasoned that inhibition of Malt1 protease activity should result in accumulation of uncleaved Malt1 substrates in these cells and allow the identification of unknown Malt1 substrates by a mass spectrometry (MS) based approach. To this end, we treated the ABC DLBCL cell line OCI-Ly3 with the Malt1 tetrapeptide inhibitor z-LVSR-fmk (Baens *et al.*, 2014) for 16h, and compared total protein levels in treated versus untreated cells, as a function of molecular weight, using a SILAC-based quantitative MS approach (Figure 1A). Inhibitor treatment led to a 3.7-fold increase in the levels of the full-length form of the previously identified Malt1 substrate RelB (Hailfinger *et al.*, 2011), validating our approach (Figure 1, B and C). Interestingly, inhibitor treatment also revealed a 5.2-fold accumulation of the full-length form of the protein Tensin-3 and a correlating decrease of a faster migrating species of Tensin-3 (Figure 1, B and C).

**Figure 1.**
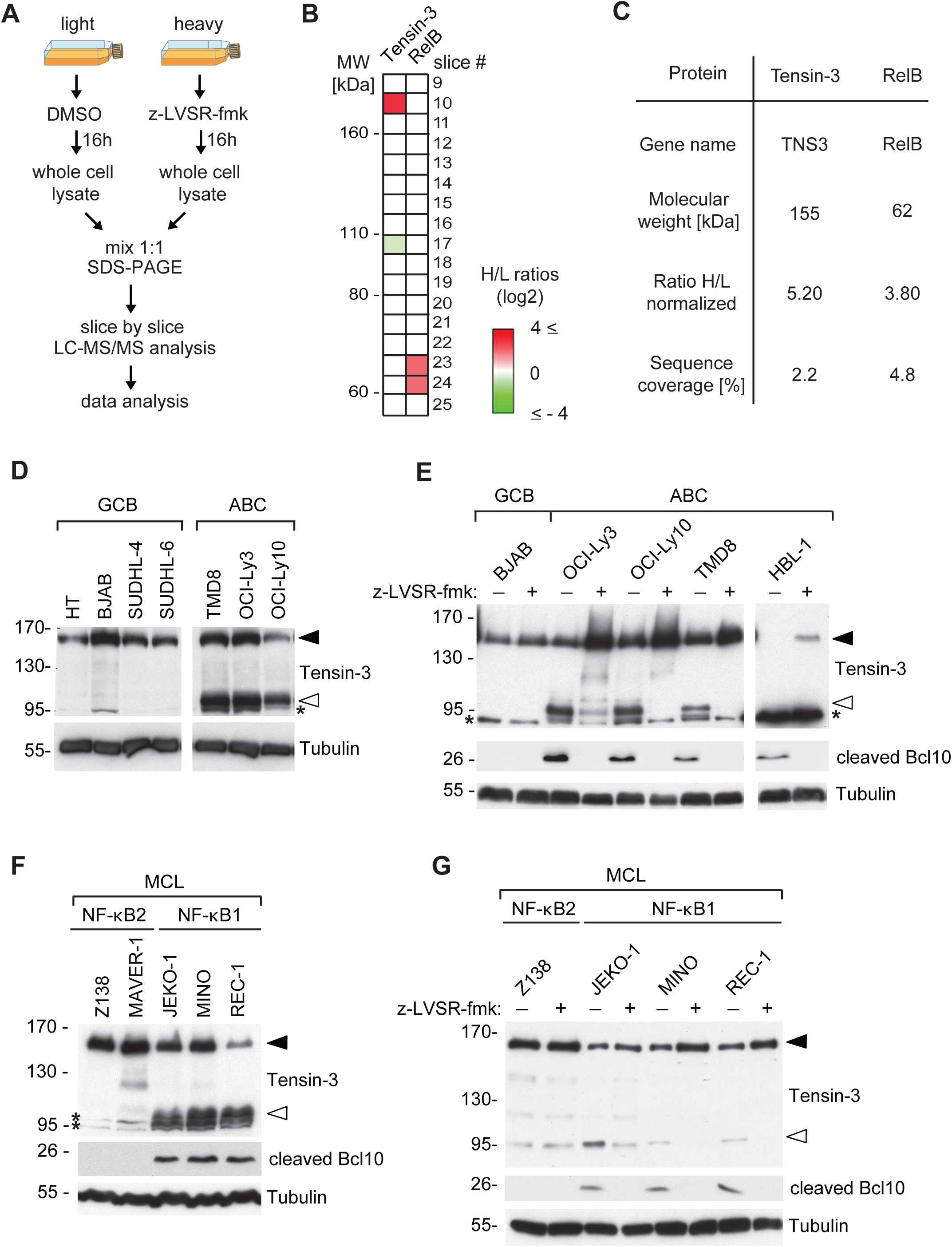
Tensin-3 is cleaved in lymphoma cell lines with constitutive Malt1 activity. (A) OCI-Ly3 cells grown in medium with “light” or “heavy” isotope-containing amino acids were treated with z-LVSR-fmk or solvent (DMSO) for 16h. Whole cell lysates were mixed, run on SDS-PAGE and the gel was cut into 51 slices that were analysed individually by tryptic digestion and liquid chromatography/tandem mass spectrometry (LC-MS/MS). (B) MS results for Tensin-3 and RelB. Color code indicates the z-LVSR-fmk-induced change in the ratio between heavy/light proteins in the indicated gel slice. (C) Table showing changes of Tensin-3 and RelB expression levels in HBL-1 cells upon treatment with z-LVSR-fmk for 16h (H, heavy isotope labelled), relative to solvent treated (L, light isotope labelled) cells. (D, E) Immunoblot analysis of the status of Tensin-3 cleavage in lysates of the indicated germinal center B cell (GCB) or activated B cell (ABC) DLBCL cell lines (D), incubated for 24h with or without z-LVSR-fmk (E). For the HBL-1 cell line, ten times more lysate was loaded for the Tensin-3 blot compared to the other cell lines. (F, G) Immunoblot analysis for Tensin-3 and cleaved Bcl10 in lysates of mantle cell lymphoma (MCL) cell lines characterized by constitutive activation of the NF-κB2 or NF-κB1 pathway (F), incubated for 24h with or without z-LVSR-fmk (G). In all figure panels, the positions of full-length and cleaved Tensin-3 is indicated by black and open arrowheads, respectively. Blotting for cleaved Bcl10 was used to monitor Malt1 activity. Tubulin serves as loading control throughout, and an asterisk indicates non-specific bands. Positions of molecular weight markers are indicated in kDa. Data are representative of a single (B, C), two (D, F, G) or three (E) independent experiments.

To validate these findings, we analysed extracts from a variety of DLBCL cell lines for the status of cleavage of Tensin-3 by Western blot, using an antibody directed against the C-terminal region of Tensin-3. This antibody detected the presence of full-length Tensin-3, migrating at the expected size of 155 kD, and a faster migrating form of Tensin-3 migrating at 95 kD, in three independent ABC DLBCL cell lines (Figure 1D). These data are consistent with the MS-based findings obtained using OCI-Ly3 cells (Figure 1, B and C). In contrast, only full-length Tensin-3 was present in GCB DLBCL cell lines such as BJAB, in which Malt1 is not active (Figure 1D). Pretreatment of the ABC DLBCL cell lines with the Malt1 inhibitor z-LVSR-fmk (Baens *et al.*, 2014) inhibited cleavage of the known Malt1 substrate Bcl10 (Rebeaud *et al.*, 2008), as expected, and led to accumulation of full-length Tensin-3 and disappearance of its faster migrating 95 kD form, suggesting that this form corresponds to a C-terminal Tensin-3 cleavage fragment (Figure 1E). In HBL-1 cells, an ABC DLBCL model that expressed low levels of Tensin-3, the cleavage fragment was below detection levels, but Malt1 inhibitor treatment also led to accumulation of full-length Tensin-3 (Figure 1E). Pretreatment with the proteasome inhibitor MG132, on the other hand, led to an increase in detection of both, N- and C-terminal Tensin-3 cleavage fragments and the previously described unstable RelB cleavage product (Figure EV1). This suggested that Tensin-3 cleavage served, at least in part, to induce its proteasomal degradation. Constitutive Malt1-dependent Tensin-3 cleavage was also detectable in a subset of mantle cell lymphoma (MCL) cell lines that are characterized by constitutive NF-κB1 activation and Malt1 activity (Rahal *et al*, 2014) (Figure 1F and 1G). In contrast, Tensin-3 cleavage was not observed in MCL cell lines relying on the NF-κB2 pathway, which lack constitutive Malt1 activity (Figure 1F and 1G).

Next, we assessed whether Tensin-3 cleavage could be induced by Malt1 activation in B-cell lines and primary B cells. In the GCB DLBCL cell line BJAB, Tensin-3 cleavage could be induced upon stimulation with the phorbolester PMA and the calcium ionophore ionomycin, which lead to activation of PKC family members and thereby mimic several aspects of BCR signaling (Figure 2A). The observed cleavage of Tensin-3, but also of another Malt1 substrate, CYLD, was efficiently inhibited by pretreatment of the cells with z-LVSR-fmk. We subsequently explored Tensin-3 expression and cleavage in primary human lymphocytes. Tensin-3 was well expressed in human primary B cells isolated from independent donors, while its levels were low to undetectable in the corresponding T cell preparations (Figure 2B). This contrasts with the expression of CYLD, which was well expressed in T-cells and, weaker, in B cells (Figure 2B). Tensin-3 levels were also very low or undetectable in various T-cell lines, including Jurkat, Hut78 and MOLT4 (not depicted). Stimulation of the primary human B cells using PMA and ionomycin led to a highly efficient induction of Tensin-3 cleavage (Figure 2C), which was blocked upon preincubation of the cells with the allosteric Malt1 inhibitor thioridazine (Nagel *et al*, 2012) (Figure 2C). Collectively, these findings identify Tensin-3 as a Malt1 substrate in activated B cells and certain B-cell lymphoma types.

**Figure 2.**
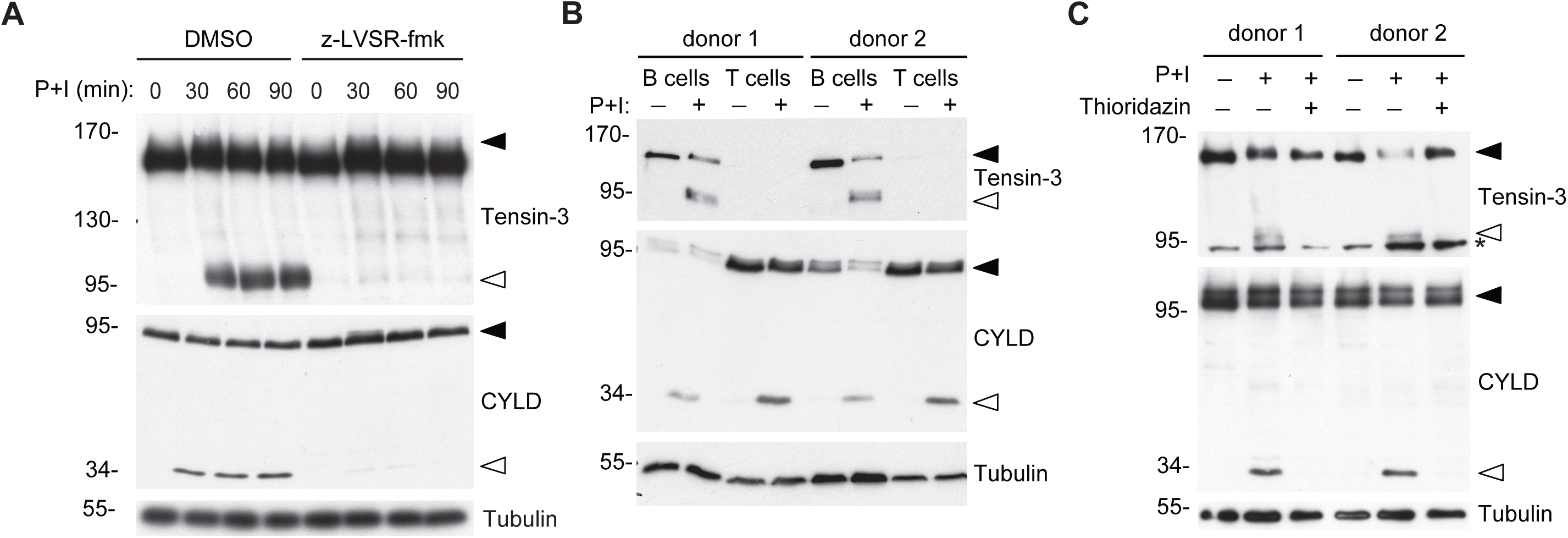
Tensin-3 is expressed in B cells and cleaved upon B-cell activation. (A) Western blot analysis of BJAB cells, pretreated with z-LVSR-fmk or solvent alone (DMSO) and stimulated for the indicated times with PMA and Ionomycin (P+I). (B, C) Immunoblot analysis of Tensin-3 cleavage in lysates of human primary CD19^+^ B and CD4^+^ T cells (B) or human primary CD19^+^ B cells (C), isolated from two independent healthy donors. Cells were incubated with or without PMA and Ionomycin (P+I) for 30 min (B, C), and pretreated with Thioridazin or solvent alone for 1h before stimulation (C). In all panels, Malt1 activity was monitored by assessing cleavage of CYLD. Black and open arrowheads indicate non-cleaved and cleaved forms, respectively, of the Malt1 substrates Tensin-3 or CYLD. Tubulin serves as loading control throughout, and an asterisk indicates non-specific bands. Data are representative of two independent experiments (A) or a single experiment, each performed with cells from two independent donors (B, C).

### Malt1 cleaves human and mouse Tensin-3 after two conserved Arg residues

To identify the Malt1-dependent cleavage site(s) in Tensin-3, we assessed its cleavage in 293T cells, in which Malt1 can be activated by its co-expression with Bcl10 (Coornaert *et al.*, 2008; Rebeaud *et al.*, 2008). Under these conditions, we observed formation of N- and C-terminal Tensin-3 fragments of approximately 72 kDa and 95 kDa, respectively (Figure 3A), which is consistent with the cleavage pattern observed for endogenous Tensin-3 (Figures 1 and 2). Cleavage was strongly impaired upon treatment of cells with the Malt1 inhibitor z-LVSR-fmk (Figure 3A). Closer analysis revealed the additional presence of less intense, faster migrating bands for both, the N- and C-terminal fragments, suggesting that Tensin-3 is cleaved at two distinct, neighboring sites (Figure 3, A and B). We therefore mutated candidate arginine residues in sequence motifs that matched the described optimal cleavage motif, Φ-x-S/P-R, in which a hydrophobic (Φ) and a variable (x) amino acid precede a Ser or Pro residue that is followed by an Arg residue, after which cleavage occurs (Hachmann *et al*, 2012). Mutation of one such Arg residue, R614 within the sequence motif LVSR_614_, strongly reduced Malt1-dependent cleavage of human Tensin-3 in 293T cells, but still allowed for residual cleavage to occur, which resulted in a C-terminal fragment with slightly lower mobility in SDS-PAGE (Figure 3B). Mutation of R645 within the sequence motif AVQR_645_ partially reduced Tensin-3 cleavage, whereas mutation of both, R614 and R645, fully prevented cleavage (Figure 3B). Interestingly, these two residues are conserved in the mouse sequence and their mutation prevented cleavage of mouse Tensin-3 upon co-expression with Malt1 and Bcl10 in 293T cells (Figure EV2). When overexpressed in BJAB cells silenced for endogenous Tensin-3 expression, we observed that stimulation with PMA and ionomycin induced cleavage of wt but not of the R614G/R645G double mutant form of human Tensin-3 (Figure 3C), even though overexpression reduced the overall efficiency of Tensin-3 cleavage when compared to the endogenous protein (see Figure 2A).

**Figure 3.**
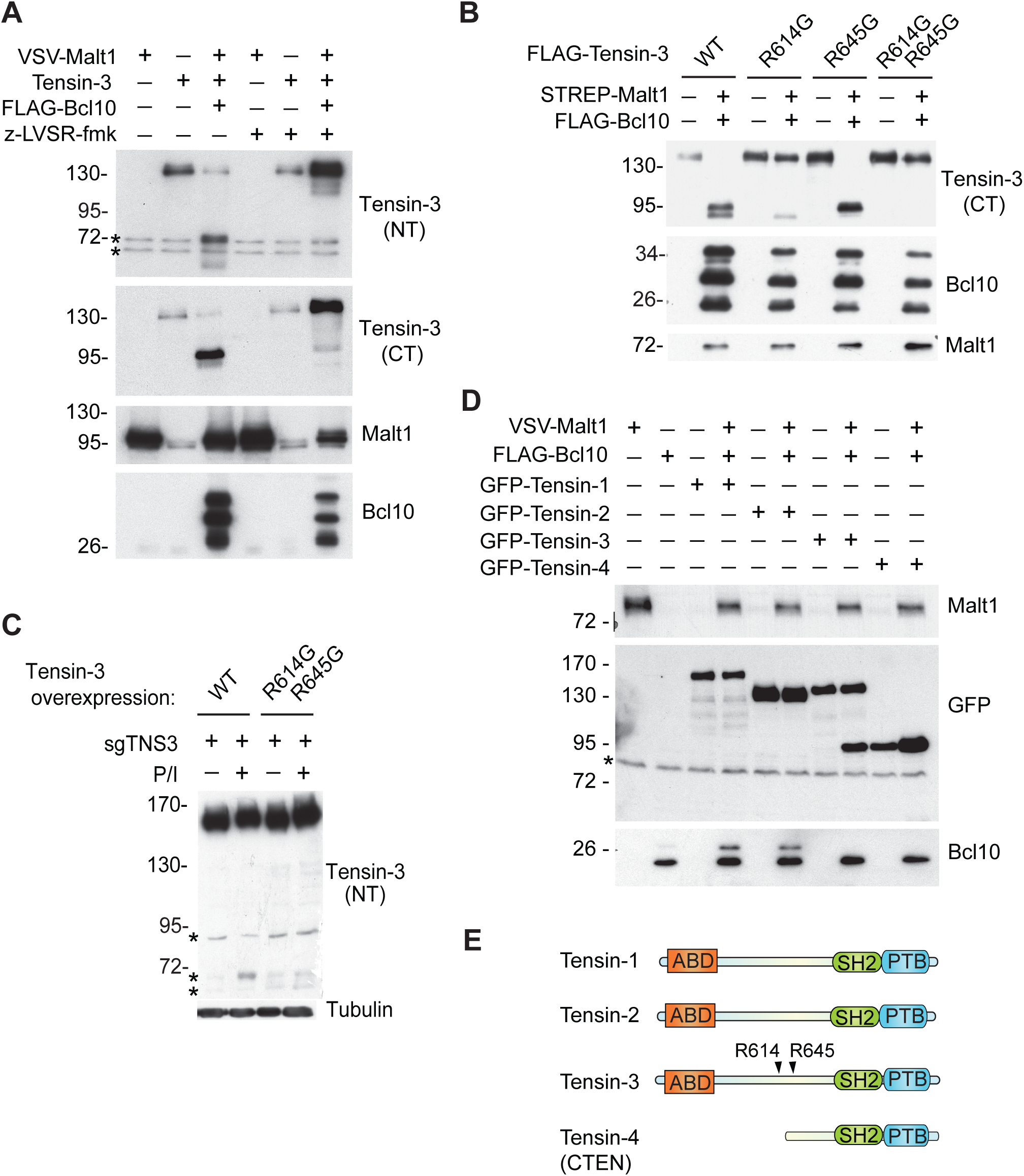
Malt1 cleaves human and mouse Tensin-3 after R614 and R645. (A) Immunoblot analysis of 293T cells transfected with the indicated combinations of VSV-tagged Malt1, untagged Tensin-3 or FLAG-tagged Bcl10, and given either no treatment (-) or treated for 16 hours (+) with z-LVSR-fmk. Tensin-3 cleavage was assessed using antibodies directed against the N-terminal (NT) or C-terminal (CT) part of Tensin-3. (B) Immunoblot analysis of 293T cells transfected with the indicated combinations of STREP-tagged Malt1, FLAG-tagged Tensin-3 or FLAG-tagged Bcl10 constructs. (C) Tensin-3-deficient BJAB cells were transduced with the indicated Tensin-3 constructs and incubated for 1 hour with (+) or without (-) PMA and ionomycin (P+I). Tensin-3 cleavage was analysed by Western blotting using an antibody directed against the N-terminal (NT) part of Tensin-3. (D) Immunoblot analysis of 293T cells transfected with the indicated combinations of GFP-tagged Tensin-1, −2, −3 and −4 (CTEN), VSV-tagged Malt1 and FLAG-tagged Bcl10. (E) Overview of the domain structure of Tensin family members. Tensin-3 is cleaved within the non-conserved sequence part, after R614 and R645. ABD: actin binding domain; SH2: src homology domain 2; PTB: phosphotyrosine binding domain. Black and open arrowheads indicate uncleaved and cleaved forms, respectively, of Tensin-3. Tubulin serves as loading control, * indicates non-specific bands. Data are representative of two (A-D) independent experiments.

Tensin-3 shares homology with Tensin-1, −2 and −4 (also known as CTEN) and the AVQR motif of the second, less efficient cleavage site of Tensin-3, is conserved in Tensin-1. Therefore, we verified whether other human Tensin proteins could be cleaved by Malt1. However, amongst the four family members, only Tensin-3 was cleaved by Malt1 (Figure 3, D and E). Thus, Malt1 activation leads to cleavage of human and mouse Tensin-3 after R614 and R645, while other Tensin family members do not seem to be targeted by Malt1.

### B-cell adhesion and -migration are impaired upon Tensin-3 silencing

So far, knowledge about the molecular function of Tensin-3 stems mainly from studies in epithelial cells, in which Tensin-3 has been reported to bind to the cytoplasmic domain of integrin β chains and to link these adhesion receptors to the actin cytoskeleton within focal adhesions (Blangy, 2017; Pylayeva & Giancotti, 2007). To assess the hypothesis that Tensin-3 plays a role in B lymphocyte adhesion, we first monitored the effect of Tensin-3 silencing on the adhesion of BJAB cells that were unstimulated or stimulated with PMA and ionomycin. Cell adhesion was analysed on plates that were coated by the β1 integrin ligand fibronectin or the β2 integrin ligands VCAM-1 or ICAM-1 (Figure 4, A and B). Deletion of Tensin-3 led to a clear reduction in the stimulation-induced cellular adhesion with all three ligands (Figure 4B). In contrast, it did not affect other, Malt1-dependent signaling events induced by stimulation, such as the phosphorylation of the NF-κB inhibitor IκB or the phosphorylation-induced activation of the JNK pathway (Figure 4C). These results obtained with Tensin-3 silenced cells suggested that Malt1-dependent Tensin-3 cleavage, which lowers the total levels of full-length Tensin-3, reduces the adhesive capacities of B cells.

**Figure 4.**
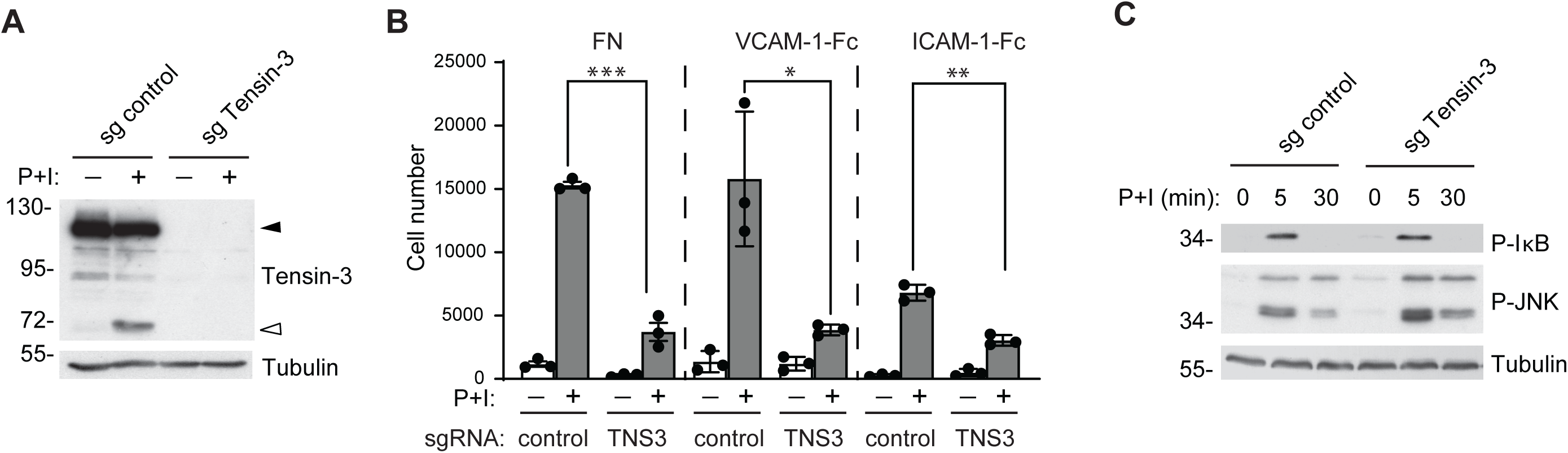
Tensin-3 levels control integrin-dependent adhesion of activated B cells. (A) Western blot analysis of BJAB cells that were transduced with Tensin-3-specific or control CRISPR/Cas9 sgRNAs and left unstimulated or stimulated with PMA and ionomycin (P+I), as indicated. (B) Adhesion of unstimulated or stimulated BJAB cells from (A) was monitored using plates coated with fibronectin (FN), VCAM-1-Fc or ICAM-1-Fc. (C) Immunoblot analysis of BJAB cells transduced with Tensin-3-specific or control CRISPR/Cas9 sgRNAs. Cells were treated with PMA and ionomycin (P+I) or solvent alone for 5 min and 30 min and lysates analysed for phosphorylation of IκBα (P-IκBα) and of JNK (P-JNK). Tubulin served as a loading control. Data are representative of two (A-C) independent experiments. Bars represent means ± SEM; differences were statistically significant with *P* < 0.05 (unpaired *t*-test) unless indicated otherwise (* P < 0.05; ** P < 0.01; *** *P* < 0.001; n.s., not significant P > 0.05).

To study the relevance of Tensin-3 cleavage in B-cell adhesion, we reconstituted Tensin-3 silenced BJAB cells with a wt or non-cleavable (nc) form of Tensin-3, mutated at both cleavage sites (Figure 5A). Unstimulated cells expressing nc-Tensin3 showed a basal increase in adhesion to fibronectin-coated plates, possibly because of low basal Malt1 activity in the cells. Compared to wt Tensin-3, cells expressing nc-Tensin3 also showed significantly increased adhesion to fibronectin upon stimulation of the cells with PMA and ionomycin (Figure 5 B). Interestingly, expression of the individual, N- or C-terminal Tensin-3 cleavage fragments did not alter the adhesive properties of BJAB cells, suggesting that cleavage abolished Tensin-3’s capacity to support adhesion (Figure 5, C and D). To assess whether Tensin-3 cleavage also played a role in the adhesion of primary B cells *in vivo*, we generated mice expressing a non-cleavable, R614A/R645A double mutant form of Tensin-3 (subsequently named TNS3-nc mice). Animals were generated by homologous recombination with a targeting vector carrying a CGC > GCC mutation (nt 816-818 of exon 17, for R614A) and an AGA > GCA mutation (nt 909-911 of exon 17, for R645A) in the Tensin-3 encoding gene Tns3 (Figure EV3A). The resulting modified Tns3 gene was distinguished from the natural Tns3 gene by PCR-based genotyping (Figure EV3B) and presence of the mutation was verified by PCR and sequencing (not depicted). Stimulation of splenic B cells with PMA and ionomycin for 30 min or 3h induced an increase in B-cell adhesiveness, and B cells expressing nc-Tensin-3 manifested a stronger capacity to adhere to fibronectin, which was statistically significant at both time points (Figure 5, E and F). Collectively, these findings support a role for human and mouse Tensin-3 in integrin-dependent B-cell adhesion and suggest that Malt1-dependent Tensin-3 cleavage serves to limit the adhesiveness of stimulated B cells, possibly to allow B-cells to detach from a cell presenting the native antigen, once BCR signaling has been triggered.

**Figure 5.**
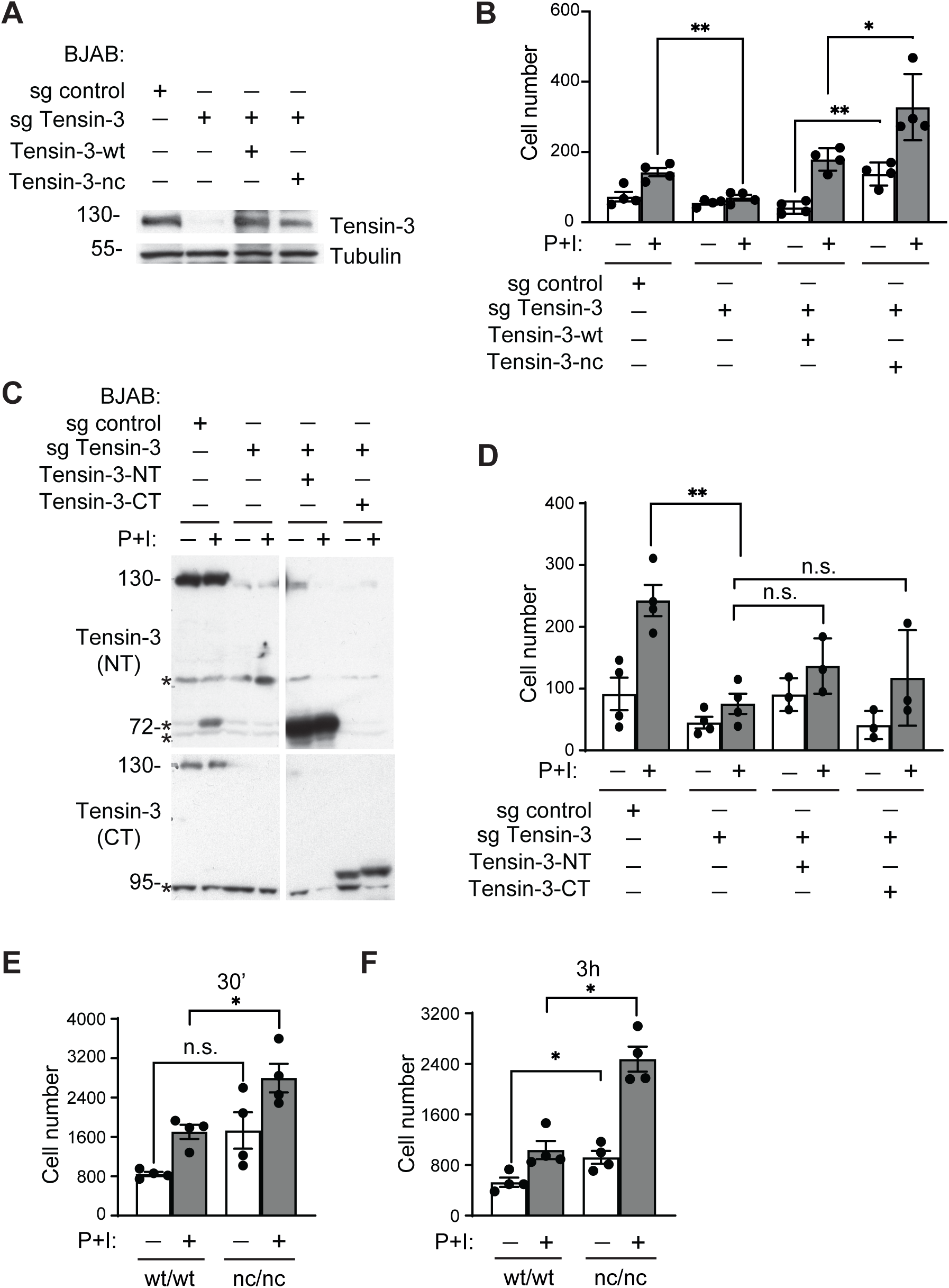
Tensin-3 cleavage limits integrin-dependent adhesion of activated B cells. (A) Immunoblot analysis of BJAB cells transduced with Tensin-3-specific or control CRISPR/Cas9 sgRNAs and reconstituted with CRISPR-resistant wt or nc Tensin-3 constructs. Tubulin served as a loading control. (B) Adhesion of BJAB cells from (C) to fibronectin-coated plates in the presence (+) or absence (-) of stimulation with PMA and ionomycin (P+I) for 30 min. (C, D) Western blot analysis (C) and adhesion to fibronectin-coated plates (D) of BJAB cells transduced with Tensin-3-specific or control CRISPR/Cas9 sgRNAs and reconstituted with the indicated, N-terminal (NT) or C-terminal (CT) Tensin-3 cleavage fragments. Cells were stimulated (+) or not (-) with PMA and ionomycin (P+I) for 30 min. (E, F) Adhesion of murine splenic B cells expressing wildtype (wt/wt) or non-cleavable Tensin-3 (nc/nc) to fibronectin-coated plates in the presence (+) or absence (-) of stimulation with PMA and ionomycin (P+I) for 30 min (E) or 3h (F). Data are representative of two (A-D) or three (E, F) independent experiments. Bars represent means ± SEM; differences were statistically significant with *P* < 0.05 (unpaired *t*-test) unless indicated otherwise (* P < 0.05; ** P < 0.01; n.s., not significant P > 0.05).

### Mice expressing a non-cleavable form of Tensin-3 show altered B cell immune responses

Based on our *in vitro* studies, we hypothesized that Tensin-3 cleavage might affect the development and/or function of B cells *in vivo*, and in particular the function of germinal center (GC) B cells, since integrin-dependent adhesion of B cells to follicular dendritic cells (FDCs) has been proposed to control the survival and fitness of GC B cells *in vitro* and *in vivo*, respectively (Koopman *et al*, 1994; Koopman *et al*, 1997; Lindhout *et al*, 1993; Wang *et al*, 2014).

Mice expressing nc-Tensin3 were born in Mendelian ratios and showed no obvious signs of developmental abnormalities (not depicted). Analysis of the absolute numbers and percentages of B- and T-cells in spleen, lymph nodes and the bone marrow revealed no significant differences between adult wild-type (wt) and TNS3-nc mice (Figure EV4, A, B and C; percentages not shown). TNS3-nc mice also had normal numbers of Treg cells in various organs, as well as follicular B cells within the spleen (Figure EV4, D and E). Marginal zone (MZ) B cells and B1a cells, which are absent or reduced in Malt1-deficient or -mutant mice (Bornancin *et al.*, 2015; Gewies *et al.*, 2014; Jaworski *et al.*, 2014; Ruefli-Brasse *et al.*, 2003; Ruland *et al.*, 2003; Yu *et al.*, 2015)), were present in normal numbers (Figure EV4, F and G).

We also tested proliferative responses of splenic B cells, isolated from naïve wildtype and TNS3-nc mice, in response to *in vitro* stimulation with anti-IgM, PMA and ionomycin, or CpG. In all settings, B cell proliferation was unaltered (Figure EV5, A-C). Next, we assessed whether Tensin-3 cleavage was required for antigen receptor-induced B cell responses *in vivo.* The basal serum levels of IgM, IgG1 (Figure 6, A), and other isotypes (Figure EV5, D) were comparable in wt and TNS3-nc mice. We next assessed whether TNS3-nc would affect the T-cell dependent humoral response by immunizing wt and TNS3-nc mice intraperitonially with NP-CGG and evaluating the splenic response 10 days later. We first noticed that total IgG1 antibodies raised against NP (NP26) were slightly reduced, while high affinity IgG1 antibodies to NP (NP4) were more strongly reduced in serum of TNS3-nc mice (Figure 6 B and C). Flow cytometric analysis of the spleen for the GC response showed that both the total and NP-specific GC B cell numbers were significantly decreased in TNS3-nc mice compared to their wt littermates (Figure 6, D, E and F). The plasma cell (PC) compartment of the spleen was also affected in mutant mice with NP-specific PC showing a reduction compared to wt counterparts (Figure 6, G and H). We thus investigated if the alteration observed in mutant mice arose mainly from a defective GC response. The ratio of dark zone (DZ, CXCR4^high^ CD86^low^) to light zone (LZ, CXCR4^low^ CD86^high^) GC B cells was unchanged (Figure 6 I). Collectively, these findings suggest that affinity maturation occurs in GC in the absence of Tensin-3 cleavage. However, PC generation and high affinity antibody responses were partially impaired.

**Figure 6.**
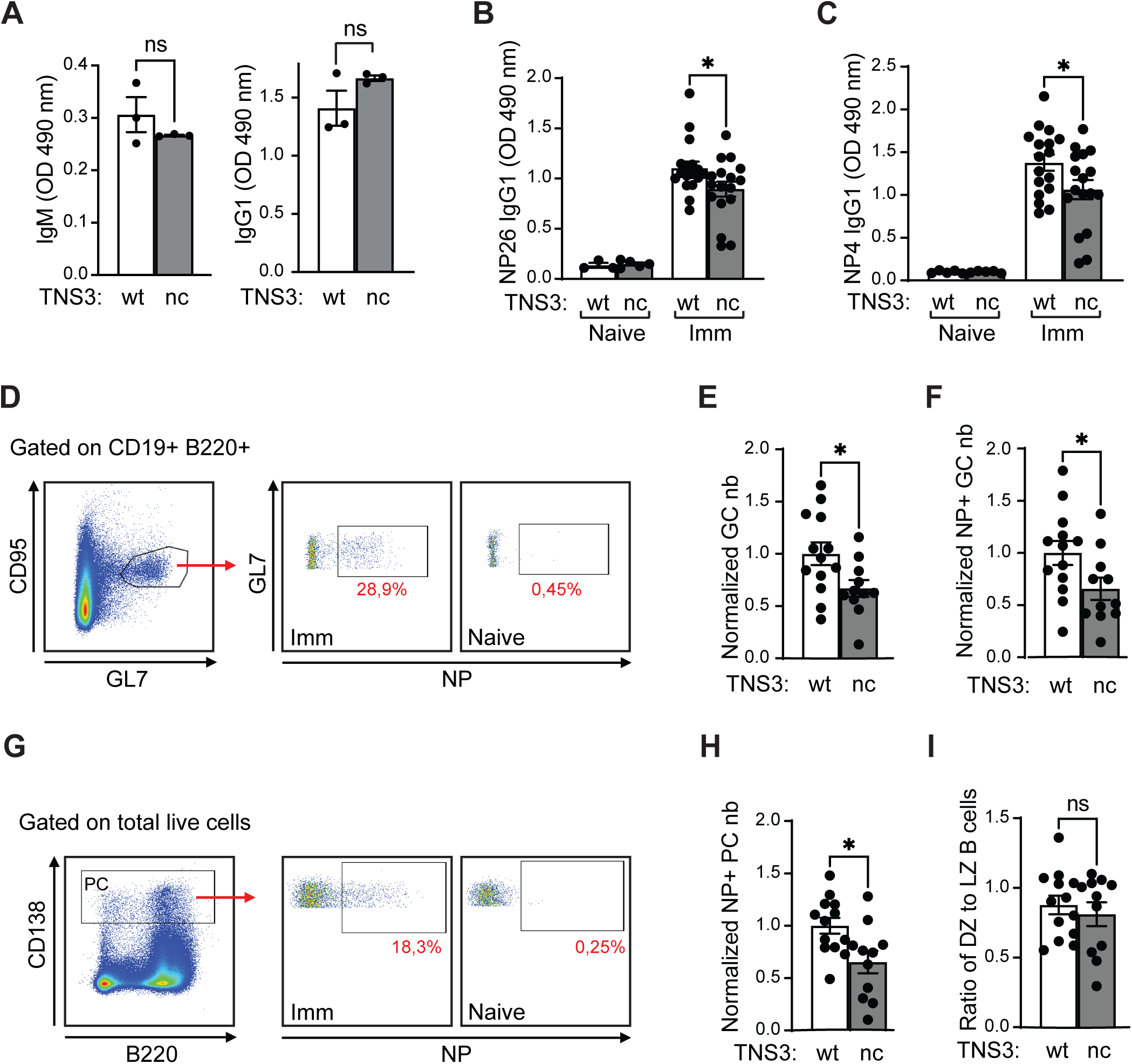
Tensin-3 cleavage is required for efficient B cell responses in vivo. (A) Basal levels of IgM and IgG1 antibodies in the serum of non-immunized wild-type (wt) and TNS3-nc mice as analysed by ELISA. (B) Analysis of wt versus TNS3-nc mice on day 10 after intraperitoneal immunization (Imm) with NP-CGG/alum versus non-immunized mice (naïve), using either ELISA for serum analysis (B, C) or flow cytometry for spleen analysis (D-I). (B, C) NP-specific antibody titers. (D) Representative dot plots showing the gating of CD95^high^ GL7^high^ GC B cells among total B cells (B220^+^ CD19^+^) (left graph) and the identification of NP-specific GC B cells in the two graphs on the right. (E, F) Quantification of the numbers (nb) of total and NP-specific GC B cells. (G) Representative dot plots showing the gating strategy to identify PC (CD138^+^) among total live cells (left graph) and the gating for NP-specific PC. (H) Quantification of NP-specific PC numbers (nb). (I) Ratio of dark zone (B220^+^ CD95^high^ GL7^high^ Cxcr4^high^ CD86^low^) versus light zone GC B cells (B220^+^ CD95^high^ GL7^high^ Cxcr4^low^ CD86^high^). Data are from one representative out of 2 (A) or a pool of 5 (B, C) or 3 (E, F, H, I) independent experiments. Bars represent means ± SEM; differences were statistically significant with *P* < 0.05 (unpaired *t*-test) unless indicated otherwise (* P < 0.05; ** P < 0.01; *** *P* < 0.001; n.s., not significant P > 0.05).

### Malt1-catalyzed Tensin-3 cleavage promotes metastasis

Finally, since we had initially identified Tensin-3 as a Malt1 substrate in ABC DLBCL models, we aimed to explore whether Tensin-3 cleavage affected the proliferation of ABC DLBCL cell lines *in vitro* and their growth and dissemination *in vivo*. Because of the large size of the Tensin-3 cDNA (4.3 kb), which greatly reduced the packaging efficiency of viral transduction constructs, we did not manage to stably express wt or nc-Tensin-3 in ABC DLBCL cell lines. Therefore, we instead focused on Tensin-3 silencing as a surrogate readout for the cleavage-dependent reduction of total Tensin-3 protein levels (see Figure 2B). First, we assessed the effect of Tensin-3 silencing on cellular growth *in vitro*, by transducing various ABC and GCB DLBCL cell lines with two different, doxycycline-inducible shRNAs that target Tensin-3. As controls, we used a non-toxic shRNA, as well as a Myc-specific shRNA that inhibits proliferation in DLBCL models. Silencing of Myc expression induced cytotoxicity in all cell lines tested (Figure 7A, **upper panel**), as previously shown (Dai *et al.*, 2017). On the contrary, silencing of Tensin-3 using two different shRNAs had no consistent effect on the growth of these cell lines *in vitro* (Figure 7A**, middle and lower panel**). We also did not observe differences in the expression of known Malt1-dependent NF-κB targets such as Bcl-XL and FLIP nor in the expression of the AP-1 family member c-Jun in sorted Tensin-3 silenced versus control cells (Figure 7B). We reasoned that, even if Tensin-3 silencing did not affect cellular proliferation of lymphoma cells *in vitro*, it might nevertheless affect the growth or dissemination of the lymphoma cells through adhesion/migration-dependent interactions with the tumor stroma and/or the neighboring cells at sites of dissemination. To test this hypothesis, we assessed the growth and dissemination of cells of the ABC DLBCL cell line HBL-1, transduced with mock or Tensin-3-targeting CRISPR/Cas9 sgRNA constructs, upon subcutaneous xenografting into the flanks of immuno-deficient NSG mice. When tumor growth was monitored over time using a caliper instrument, we noticed no significant differences in tumor growth and tumor weight between the control and Tensin-3-deficient cells (Figure 7C, D). However, when we analysed the bone marrow for the presence of infiltrating hCD20^+^ tumor cells, we detected a significantly enhanced dissemination of Tensin-3 silenced cells compared to control cells (Figure 7E). Together, these findings suggest that Malt1-dependent Tensin-3 cleavage, which lowers cellular Tensin-3 levels, enables cells to detach from the tumor site and spread to a distant location.

**Figure 7.**
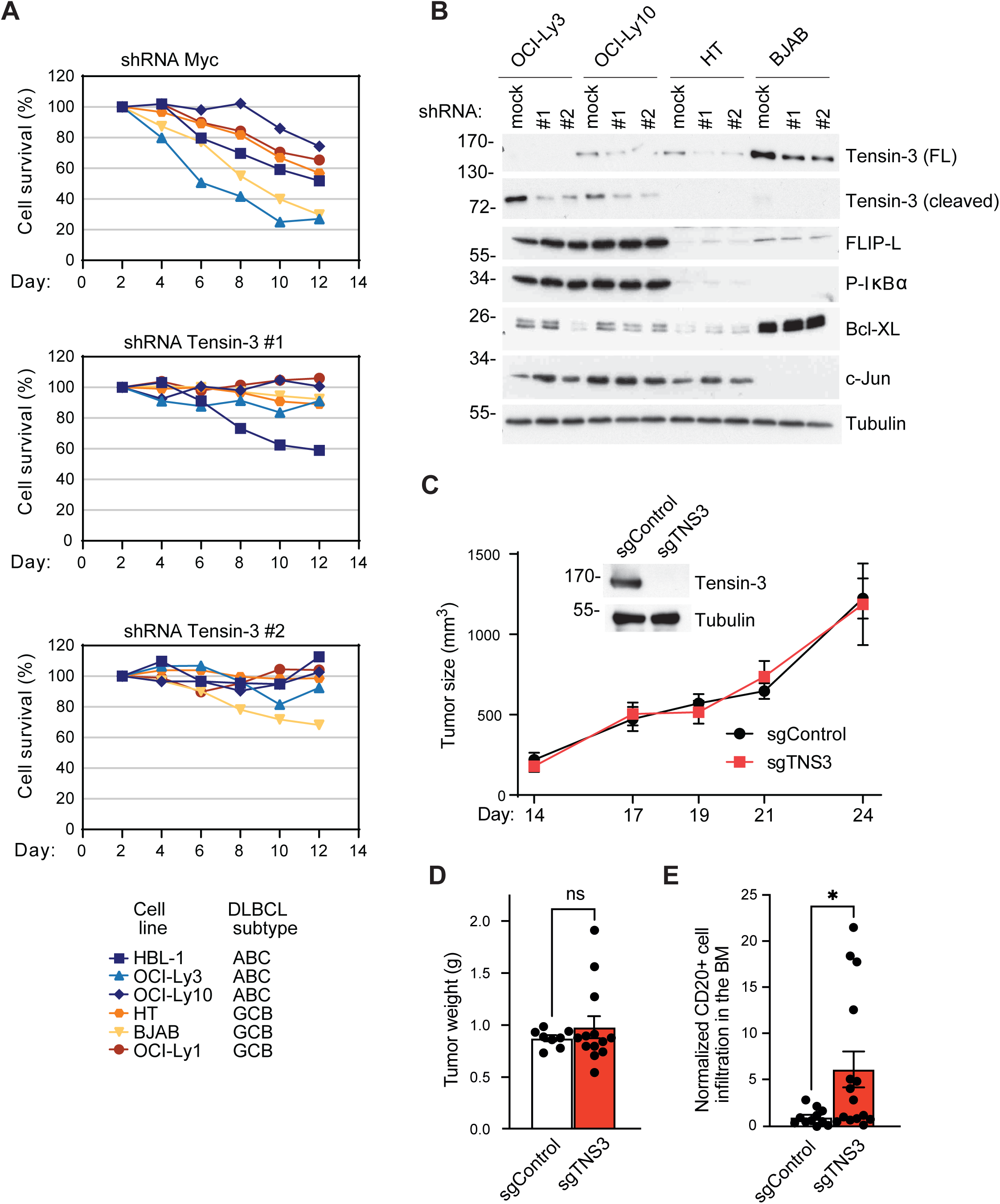
Role of Tensin-3 in the dissemination of xenografted ABC DLBCL cells. (A) DLBCL cell lines of the ABC (HBL1, OCI-Ly3, OCI-Ly10) or GCB subtype (HT, BJAB, OCI-Ly1) were transduced with Myc- or Tensin-3-specific shRNAs and viability of transduced, GFP-positive cells was monitored over time upon inducible shRNA expression. (B) Western blot analysis of the cell lines used in (A), 3 days after induction of the two different Tensin-3-specific shRNAs. Positions of molecular weight markers are indicated on the left. (C, D) Tumor growth of xenografted HBL-1 cells stably transduced with control or Tensin-3-specific sgRNA (C) and weight of tumors excised from mice at day 24 (D). (E) Flow cytometric analysis of the presence of hCD20+ HBL-1 cells used in (C) to the bone marrow of NSG mice 24 days after xenografting of tumor cells into the flanks of the animals. Data are representative of two (A-C) independent experiments. (D) and (E) show pooled data of 2 independent experiments. In (E), data were normalized to control samples. Bars represent means ± SEM; differences were statistically significant with *P* < 0.05 (unpaired *t*-test) unless indicated otherwise (* P < 0.05; ** P < 0.01; *** *P* < 0.001; n.s., not significant P > 0.05).

## Discussion

Here, we identify the scaffold protein Tensin-3 as a novel Malt1 substrate and provide several lines of evidence for a role of Tensin-3 in B-cell adhesion that is negatively regulated by its Malt1-dependent cleavage. First, full-length Tensin-3 levels were greatly reduced in activated human B cells and in lymphoma cell lines characterized by constitutive Malt1 activity, giving rise to cleaved Tensin-3 fragments. Second, human B cells lacking Tensin-3 had an impaired capacity for integrin β1 and β2-dependent adhesion, whereas adhesion was strengthened in human and mouse B cells expressing a non-cleavable compared to a wt form of Tensin-3. Third, Tensin-3 cleavage was required for an efficient germinal center reaction, which relies on integrin-dependent B-cell adhesion to FDCs. Finally, we observed that silencing of Tensin-3 levels promoted the metastatic spreading of xenografted lymphoma cells. Collectively, these findings support a model in which Malt1-dependent Tensin-3 cleavage downregulates the adhesive properties of activated B cells and lymphoma cells.

The growing family of cellular Malt1 substrates includes Bcl10 (Rebeaud *et al.*, 2008), RelB (Hailfinger *et al.*, 2011), A20 (Coornaert *et al.*, 2008), HOIL-1 (Douanne *et al.*, 2016; Elton *et al.*, 2016; Klein *et al.*, 2015), CYLD (Staal *et al.*, 2011), Regnase-1 (Uehata *et al.*, 2013), Roquin (Jeltsch *et al.*, 2014) and N4BP-1 (Yamasoba *et al.*, 2019), and Malt1 itself (Baens *et al.*, 2014). Two additional proteins, the kinase NIK and the tumor suppressor LIMA1, are cleaved exclusively by the oncogenic API2-Malt1 fusion protein (Nie *et al.*, 2015; Rosebeck *et al*, 2011). Most of the presently known Malt1 substrates regulate gene transcription during inflammatory immune responses. Tensin-3 is unique amongst the presently known Malt1 substrates for two reasons: first, it is not expressed in mature T cells and second, it controls a transcription-independent aspect of the immune response by specifically regulating the integrin-mediated adhesiveness of B cells. Interestingly, two other Malt1-substrates, both with previously described roles in transcriptional regulation, have been proposed to have additional roles in cellular adhesion: Bcl10, whose cleavage promotes T-cell adhesion by β1 integrins through unknown mechanisms (Rebeaud et al., 2008), and CYLD, whose cleavage induces disruption of microtubules and a correlating loss of adhesiveness of vascular endothelial cells (Klei *et al.*, 2016). Thus Tensin-3 is part of an emerging group of Malt1 substrates whose cleavage controls transcription-independent aspects of immune responses, by regulating cellular adhesiveness.

In the past, the role of Tensin-3 has been explored exclusively in non-immune cells. These studies have supported the idea that Tensin-3 acts as a negative regulator of cell migration and growth in various untransformed and cancer cell lines (Chen *et al*, 2017; Katz *et al*, 2007; Martuszewska *et al*, 2009; Pylayeva & Giancotti, 2007). Interestingly, Tensin-3 levels have been shown to be downregulated in tumor cells by various mechanisms, including inhibition of TNS3 gene transcription (Katz *et al.*, 2007) and Tensin-3 mRNA translation (Chen *et al.*, 2017). Moreover, EGFR triggering can induce recruitment and phosphorylation of Tensin-3, resulting in its release from focal adhesion proteins (Cui *et al*, 2004). Overall, these mechanisms all downregulate or sequester Tensin-3 with the apparent aim to reduce cellular adhesion and promote the migration and invasive spreading of carcinoma cells. Another interesting way of Tensin-3 regulation is through an inducible switch in expression from Tensin-3 towards expression of the Tensin homologue CTEN, whose structure closely matches the Malt1-induced C-terminal Tensin-3 fragment. This Tensin-3/CTEN switch has been proposed to promote cellular migration by displacing Tensin-3 from beta1 integrins, thereby disassembling fibrillar cell contacts (Katz *et al.*, 2007; Pylayeva & Giancotti, 2007). In view of these findings, it is tempting to speculate that Malt1-dependent Tensin-3 cleavage might, at least in part, promote migration of B cells by lowering the levels of Tensin-3 and generating a CTEN-like Tensin-3 cleavage fragment that would compete with full length Tensin-3 for the binding to the integrin beta chain.

Triggering of the antigen receptor on B cells induces a rapid increase in integrin-mediated cellular adhesion (Dang & Rock, 1991; Koopman *et al*, 1991; Spaargaren *et al*, 2003). We propose that BCR-induced Malt1 activation, which builds up slowly and reaches a peak after approximately 30 min of stimulation (Rebeaud *et al.*, 2008), will limit the duration of B-cell adhesiveness by inducing cleavage and subsequent degradation of Tensin-3, which should loosen the anchoring of beta-integrins to the cellular actin cytoskeleton (Pylayeva & Giancotti, 2007). B cell adhesion plays an important role in the interaction of B cells with FDCs that display native antigens in the form of immune complexes (Kosco-Vilbois, 2003). α4β1 and αLβ2 (LFA1) integrins were shown to be involved in this interaction and to contribute to the proliferation and survival of GC B cells *in vitro* (Koopman *et al.*, 1994; Koopman *et al.*, 1997; Lindhout *et al.*, 1993). *In vivo* studies, however, have suggested an alternative role for integrin-mediated interactions of FDCs with GC B cells in regulating the fitness rather than the survival of GC B cells (Wang *et al.*, 2014). In our hands, activated B cells isolated from Tensin-3-nc mice showed an increase in the intensity and duration of integrin-dependent adhesiveness compared to wild-type B cells *in vitro*. Moreover, the mice expressing a non-cleavable form of Tensin-3 displayed a reduced high affinity antibody response upon immunization, which affected to a similar extent the germinal center response and the PC numbers and led to reduced high affinity titers of NP-specific IgG1 antibodies. Together, these observations point to a specific defect in GC B cell responses of Tensin-3-nc mice, presumably due to a slower detachment of naïve B cells from FDCs once they have captured native antigen.

While Tensin-3-nc mice showed partially reduced humoral immune responses, we did not observe any of the other developmental and functional immune defects reported for mice expressing a catalytically inactive form of Malt1, such as impaired development of MZ and B1 B cells, reduced Treg numbers and impaired T-cell responses (Bornancin *et al.*, 2015; Gewies *et al.*, 2014; Jaworski *et al.*, 2014). The lack of a T-cell phenotype correlates with our observation that Tensin-3 expression was not detectable in mature primary T cells and T-cell lines. Moreover, the development of MZ and B1 B cells and of Treg cells relies to a large extent on the canonical NF-κB response (Gerondakis *et al*, 2006; Zhang et al, 2017), which is driven by Malt1-dependent cleavage of RelB and A20 and by Malt1 autoprocessing (Baens *et al.*, 2014; Coornaert *et al.*, 2008; Hailfinger *et al.*, 2011) but independent of Tensin-3. The only noticeable phenotype of the Tensin-3 nc mice was a defect in the T-cell dependent humoral immune response, which is also impaired or delayed in mice expressing catalytically inactive Malt1 (Bornancin *et al.*, 2015; Gewies *et al.*, 2014; Jaworski *et al.*, 2014; Yu *et al.*, 2015) and thus likely to rely, at least in part, on Malt1-dependent Tensin-3 cleavage.

Finally, our study provides new insight into the mechanism underlying the extranodal dissemination of DLBCL cells with constitutive Malt1 activity. ABC subtype-typical mutations affecting *MyD88* and/or *CD79B* are more frequent in patients with multiple extranodal involvements (ENIs) compared to patients with no or single ENI (Shen *et al*, 2020). Amongst newly diagnosed patients with DLBCL, approximately 10% have bone marrow infiltrates, and the presence of concordant bone marrow involvement is associated with poorer outcome (Campbell *et al*, 2006; Chung *et al*, 2007; Muringampurath-John *et al*, 2012; Sehn *et al*, 2011). When assessing the bone marrow dissemination of xenografted ABC DLBCL HBL-1 cells, which harbor a *MyD88* and a *CD79B* mutation, we observed that silencing of Tensin-3 promoted dissemination of lymphoma cells to the bone marrow, without affecting tumor growth by itself. These findings suggest that Malt1-dependent cleavage of Tensin-3 may contribute to the spreading of ABC DLBCL to distant sites, possibly by promoting the detachment of tumor cells from FDCs or other cells resident in B cell follicles at the site of the primary tumor that typically arises at a lymph node.

In conclusion, our work reveals a previously unexpected contribution of Tensin-3 to the immune response and lymphomagenesis. Moreover, our findings suggest that Malt1 controls specific aspects of the humoral immune response and the dissemination of lymphoma cells in a transcription-independent manner, by modulating cellular adhesion through Tensin-3 cleavage.

## Expanded figure legends

**Expanded Figure EV1.**
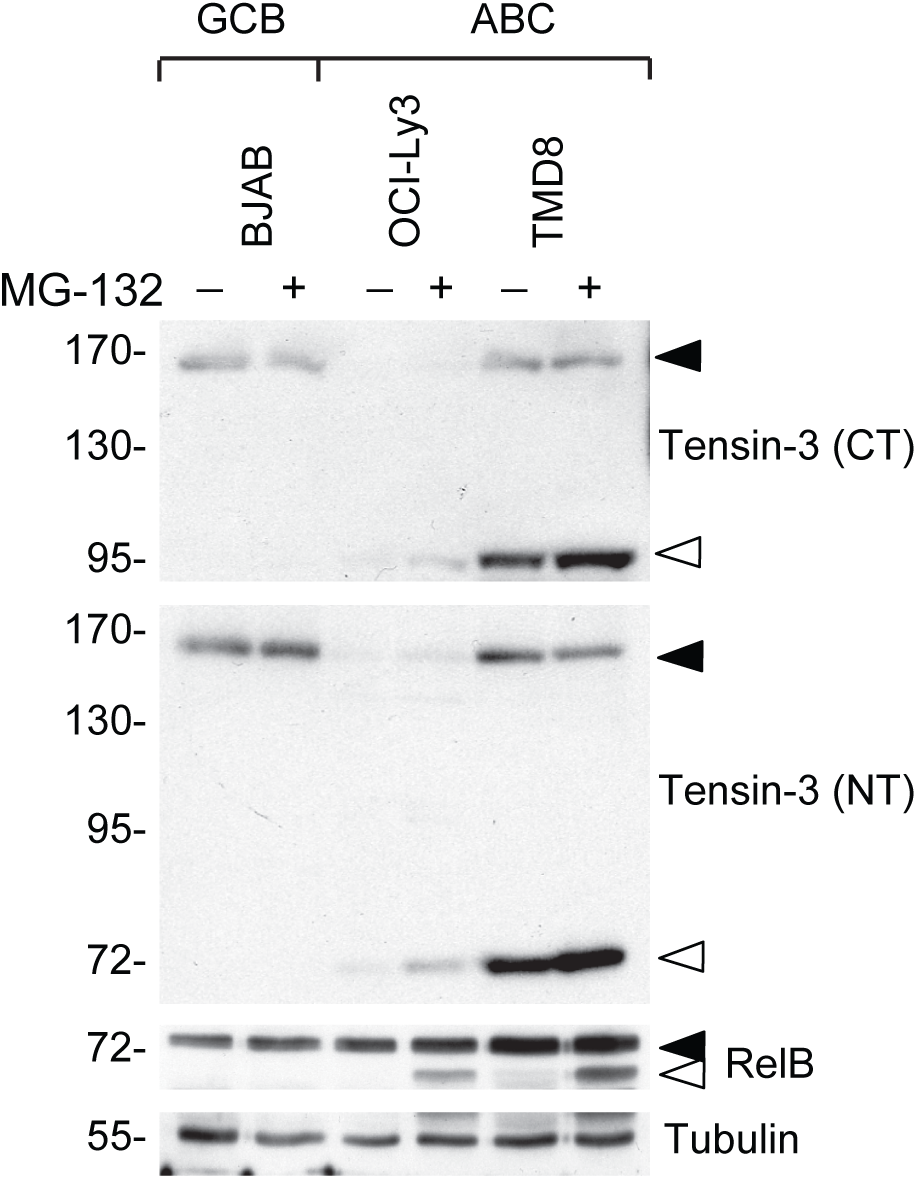
Proteasomal inhibition stabilizes Tensin-3 cleavage fragments. Immunoblot analysis of Tensin-3 and RelB cleavage in lysates of the indicated DLBCL cell lines treated with MG-132 (+) or solvent (DMSO) (-). Tensin-3 cleavage was assessed using antibodies directed against the C-terminal (CT) or N-terminal (NT) part of Tensin-3, and efficiency of MG-132 treatment by monitoring stabilization of the RelB cleavage product. Tubulin served as a loading control. Data are representative of two independent experiments.

**Expanded Figure EV2.**
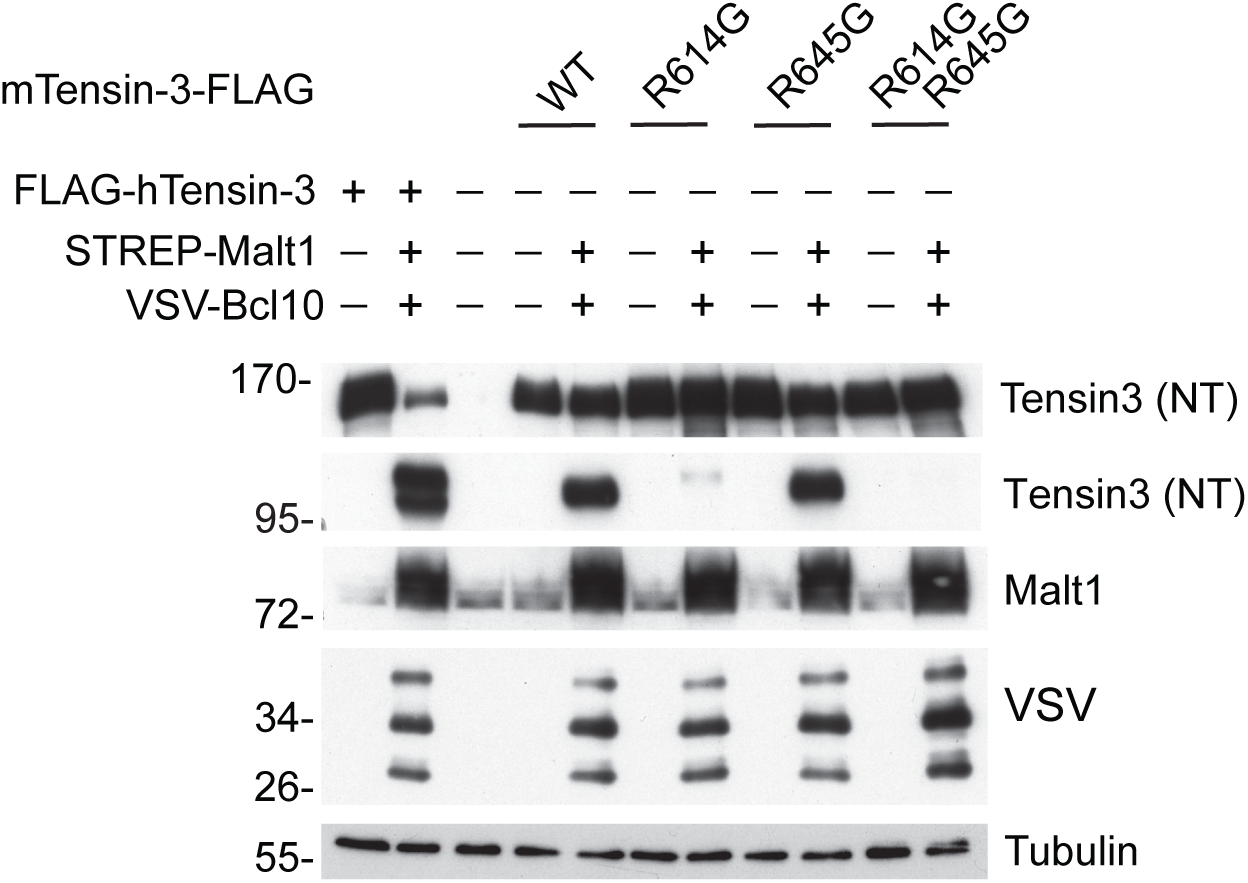
Malt1 cleaves mouse Tensin-3 after R614 and R645. Immunoblot analysis of 293T cells transfected with the indicated combinations of STREP-tagged human Malt1, N-terminally FLAG-tagged human Tensin-3 (hTensin-3), C-terminally FLAG-tagged mouse Tensin-3 (mTensin-3) or VSV-tagged human Bcl10. Data are representative of two independent experiments.

**Expanded Figure EV3.**
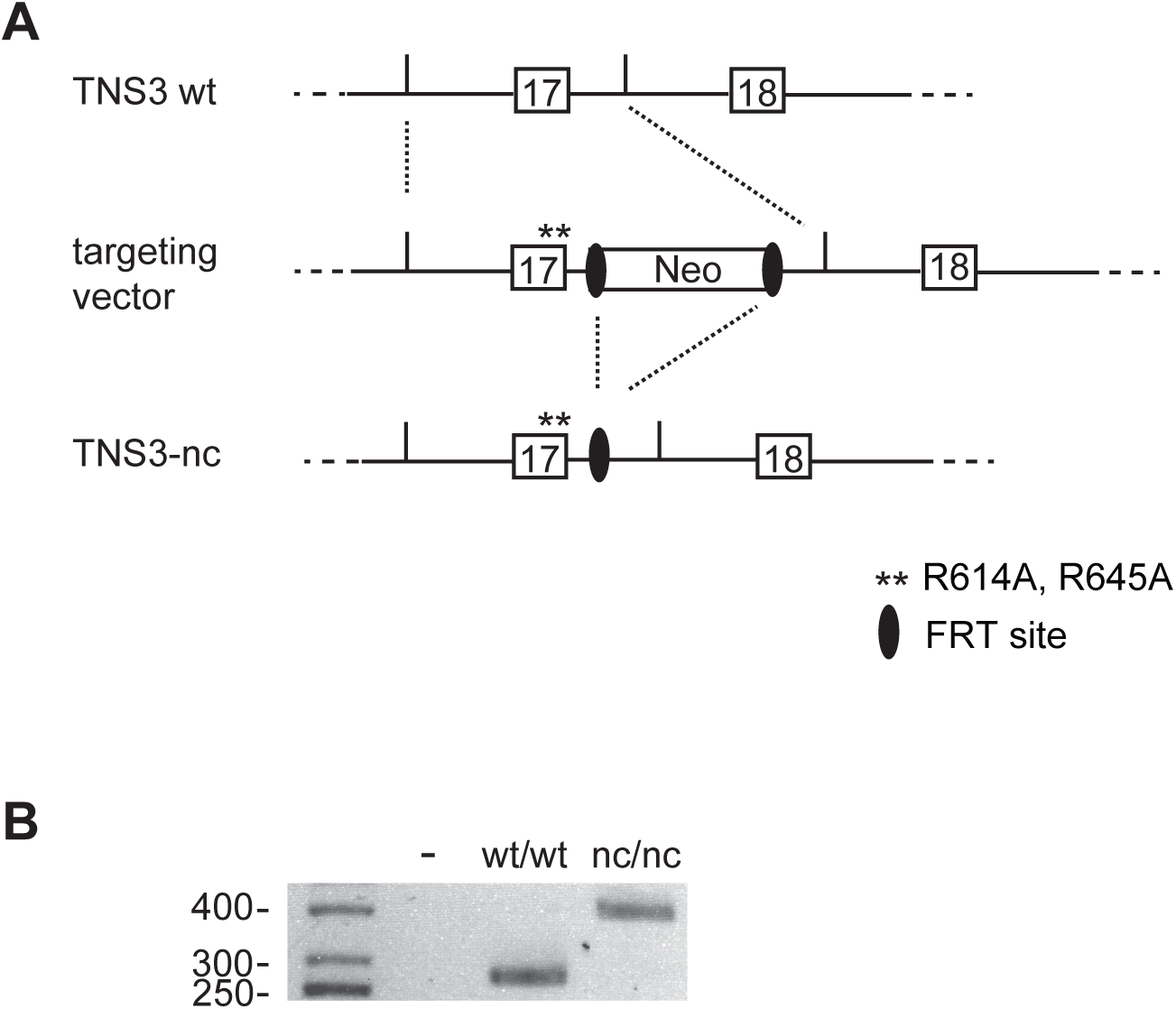
Strategy used to generate and analyze TNS3-nc mice. (A) Overview of the targeting vector used to generate mice expressing non-cleavable (nc) Tensin-3. The vector carries two point-mutations in exon 17, introducing the following sequence alterations: CGC>GCC for R614A and AGA > GCA for R645A, and a neomycin resistance cassette (Neo) flanked by flippase recombinase target (FRT) sites. In the Tensin-3 non-cleavable (TNS3-nc) mice used for this study, the neomycin resistance cassette was removed by crossing to mice expressing the flippase recombinase (FLP). (B) Genotyping of homozygous wild-type (wt/wt) and TNS3-nc (nc/nc) mice by PCR amplification of the FRT region, generating a 391 or 277 bp fragment, respectively. Position of molecular weight markers (in bp) is indicated.

**Expanded Figure EV4.**
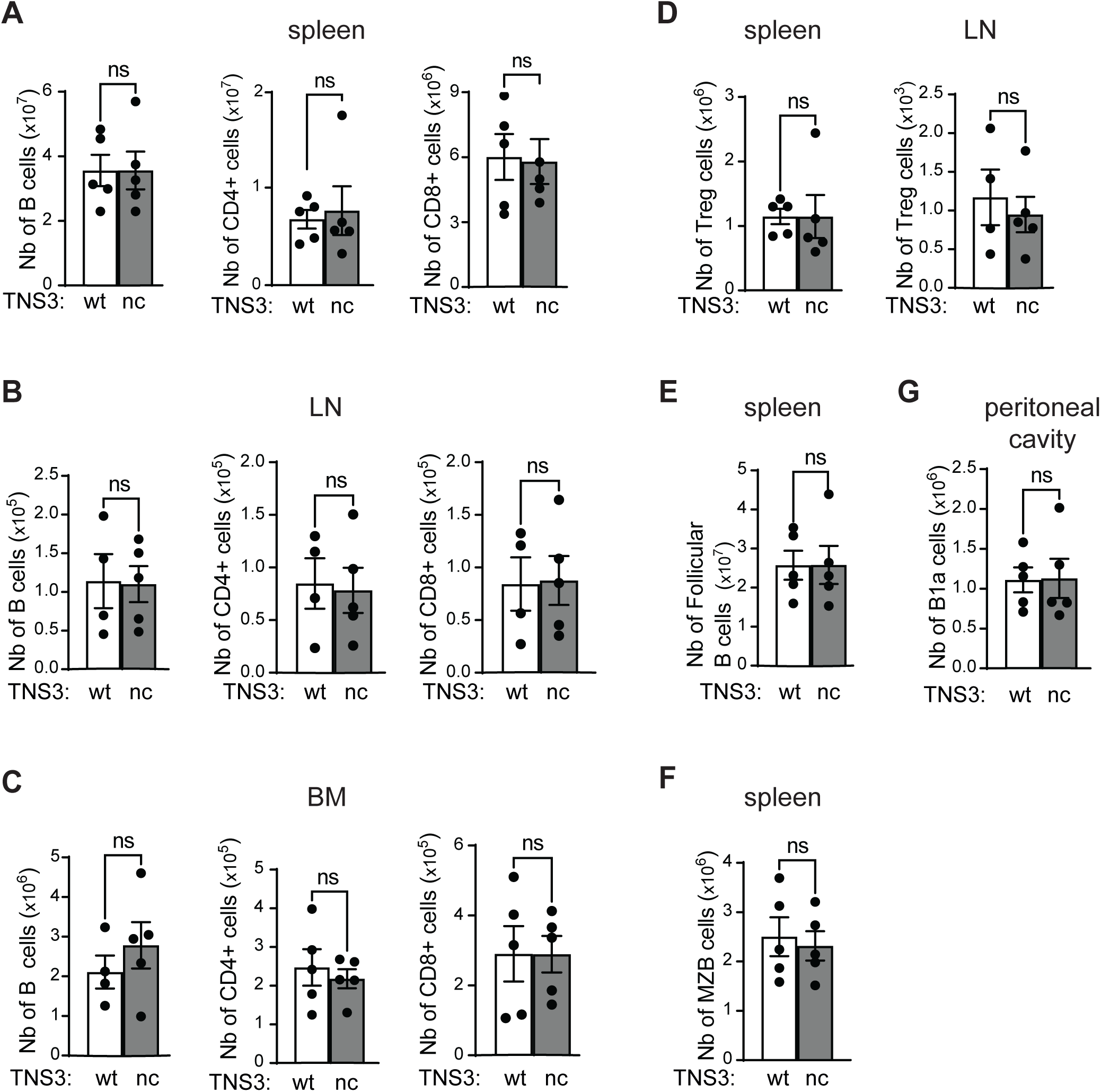
Analysis of B- and T-cell subsets in wt and TNS3-nc mice. Flow cytometric analysis of immune cell types in mice expressing non-cleavable Tensin-3 (TNS3-nc) and wild-type (wt) mice, plotted as absolute cell number in a given organ. (A-C) B cells (CD19+B220+) and CD4^+^ or CD8α^+^ T cells in the spleen (A), inguinal lymph nodes (B) and bone marrow from one femur and one tibia (C). (D) CD4^+^ FoxP3^+^ Treg cells in spleen and lymph nodes (LN). (E, F, G) CD19^+^ B220^+^ CD23^+^ CD21^-^ follicular B cells (E) and CD19^+^ B220^+^ CD23^-^ CD21^+^ marginal zone B (MZB) cells (F) in the spleen, and of CD19^+^ CD5^+^ B1a (G) cells in the peritoneal cavity. Data are representative of 2 independent experiments. Bars represent means ± SEM; differences were statistically significant with *P* < 0.05 (unpaired *t*-test) unless indicated otherwise (* P < 0.05; ** P < 0.01; *** *P* < 0.001; n.s., not significant P > 0.05).

**Expanded Figure EV5.**
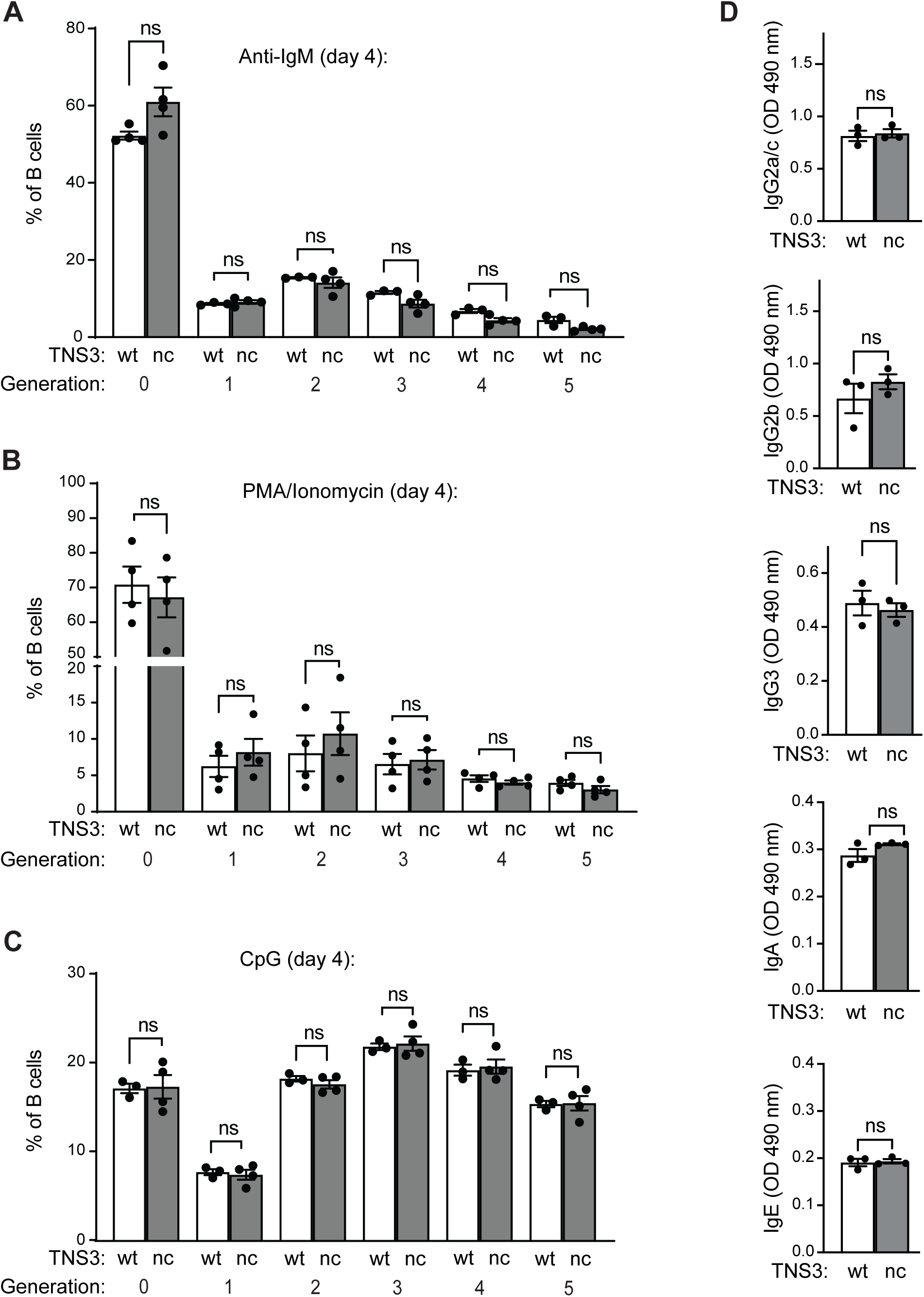
Analysis of B-cell responses in wt and TNS3-nc mice. (A-C) Proliferation of CTV-labelled CD19^+^ B220^+^ cells from wild-type (wt) and TNS3-nc mice in response to *in vitro* stimulation with anti-IgM (A), PMA and ionomycin (PI) (B) or CpG (C). (D) Basal levels of IgG2a/c, IgG2b, IgG3, IgA and IgE in the serum of non-immunized wild-type (wt) and TNS3-nc mice. Data are from one representative out of 3 (A-C) and from 2 (D). Bars represent means ± SEM; differences were statistically significant with *P* < 0.05 (unpaired *t*-test) unless indicated otherwise (* P < 0.05; ** P < 0.01; *** *P* < 0.001; n.s., not significant P > 0.05).

## Materials and methods

### Identification of Malt1 substrates by quantitative proteomics

The identification of Malt1 substrates by SILAC, protein separation and quantitative MS analysis was essentially performed as described before (Di Micco *et al*, 2016; Morikawa *et al*, 2014). Briefly, OCI-LY-3 cells were cultured in modified RPMI medium containing either heavy isotope-labelled or unlabelled Lysine and Arginine for 6 passages to achieve complete labelling (>98%). Cells cultured in heavy or light medium were treated for 16h with z-LVSR-fmk (2 μM) or solvent alone (DMSO), respectively. Cell lysates were mixed at a quantitative ratio of 1:1, and 120 ug of protein were resolved on a gradient SDS-PAGE A total of 120 μg of protein from mixed lysates was migrated on a pre-cast 4-12% Novex NuPAGE Bis-Tris SDS Mini Gel (Invitrogen). After Coomassie staining, the lane was divided into 51 individual horizontal gel slices, followed by in gel tryptic digestion and the resulting peptides in the supernatant were concentrated and analysed by LC-MS/MS on a LTQ-Orbitrap Velos mass spectrometer (Thermo Fisher Scientific) as described (Morikawa *et al.*, 2014). Protein identification in gel slices was performed with MaxQuant 1.3.0.5 (Cox & Mann, 2008). Proteins with different normalized SILAC ratios were visualized as a function of the gel slice using custom made scripts as previously described (Di Micco et al., 2016).

### Antibodies

Antibodies used included polyclonal rabbit anti-human Tensin-3 (C2, Santa Cruz, directed against the C-terminus), affinity purified rabbit anti-human Tensin-3 (generated against a GST-Tensin-3 fusion protein comprising amino acids 418–577 of human Tensin-3), rabbit anti-Bcl10 (H197, Santa Cruz Biotechnology), anti-Tubulin, anti-FLAG (M2), anti-VSV (polyclonal and monoclonal P5D4) and anti-pERK (MAPK-YT) (Sigma), anti-pIκBα, anti-IκBα, anti-RelB and anti-CYLD (Cell Signaling), anti-Strep-HRP (IBA BioTAGnology) and anti-GFP (Enzo LifeSciences). Affinity-purified Malt1 antibodies and antibodies specific for cleaved Bcl10 have been previously described (Hailfinger *et al.*, 2009; Rebeaud *et al.*, 2008). Horseradish peroxidase–coupled goat anti-mouse or anti-rabbit were from Jackson Immunoresearch. Anti-human CD20-FITC (2H7) and FoxP3 PE (FJK-16s) for flow cytometry were purchased from eBioscience. Anti-mouse antibodies for flow cytometry included CD19 PE-TexasRed (1D3), B220 BV421 (RA3-6B2), CD138 BB515 (281-2), CD23 BB700 (B3 B4), CD8α APC-H7 (53-6.7) were purchased from BD Biosciences. Anti-mouse CD21 PE-Cy-7 (eBioBD9), IgM APC (II/41), CD5 APC (53-73) were purchased from eBioscience. Anti-mouse Cxcr4 PE (2811) was from Invitrogen. Anti-mouse CD86 FITC (GL-1), B220 AF700 (RA3-6B2), GL7 Pacific blue (GL7), CD95 PerCp Cy5.5 (SA367H8), TCRβ APC-Cy7 (H57-597) were from Biolegend. Anti-mouse CD4 PerCP (RM 4-5) was from BD Pharmingen. NP-PE was purchased from Biosearch Technologies.

### Cell culture and cell stimulation

HEK293T cells were cultured in DMEM supplemented with 10% FCS and antibiotics, respectively. Lentivirally transduced cells were always kept under puromycin selection (5 µg/ml). The diffuse large B cell lymphoma cell lines BJAB, SUDHL-4, SUDHL-6, HT, OCI-Ly1, HBL-1, OCI-Ly3, OCI-Ly10, TMD8 and U2932 and the mantle cell lymphoma cell lines Z138, MAVER-1, JECO-1, MINO, REC-1 were cultured as described (Hailfinger *et al.*, 2009; Juilland *et al*, 2016) and their identity was certified by genotype-based cell line authentication (Microsynth). To stimulate B- and T cells, we used PMA (10 ng/mL; Alexis) and ionomycin (1 μM; Calbiochem). In some experiments, cells were preincubated with z-LVSR-fmk (2 μM, Bachem), Thioridazine (10 μM, Sigma) or MG132 (5 uM, Calbiochem).

### Plasmids

Expression constructs for GPF-Tensin-1, GFP-Tensin-2, GFP-Tensin-3 and GFP-Tensin-4 were a kind gift from Katherine Clark (University of Leicester). The lentiCRISPRv2 vector (GeCKO) was used for CRISPR/Cas9-mediated Tensin-3 silencing. Reconstitution with Tensin-3 expression constructs was performed using a previously described retroviral vector (Neal & Clipstone, 2002). Silent point mutations and non-cleavable point mutants were generated by quick-change PCR using Kapa high-fidelity DNA polymerase (Roche) and all mutations were verified by sequencing.

### Transfection and transduction of cells

Transient transfection of HEK293T cells and lentiviral transduction of lymphocyte cell lines were essentially performed as previously described (Rebeaud *et al.*, 2008). To stably silence Tensin-3 expression, BJAB and HBL-1 cells were stably transduced with two Tensin-3-specific single guide RNAs (sgRNA) (5’-GTACAGTGGGACCCGCCACG -3’, 5’-GTGTACCATATGCAAGGCGC-3’) or control (luciferase-specific) sgRNA (5’-CTTCGAAATGTCCGTTCGGT-3’) and selected using puromycin. Cells were subsequently transduced to constitutively express GFP together with wt or non-cleavable Tensin-3 constructs that were rendered CRISPR/Cas9 resistant by a silent point mutation. Transduced cells were sorted for live GFP positive cells using flow cytometry. The shRNA-mediated RNA interference and cytotoxicity assays were performed as previously described (Nogai *et al*, 2013). DLBCL cell lines were transduced with two Tensin-3 shRNA (5’-CCACTGTTTGCCATCATCTAA-3’ and 5’ GTTCTGGTACAAGGCGGATAT-3’) or a Myc shRNA (‘CGATTCCTTCTAACAGAAATG’).

### Lysis, immunoprecipitation and immunoblotting

Cells were lysed in lysis buffer containing 50 mM HEPES pH 7.5, 150 mM NaCl, 1% Triton-X-100, protease inhibitors (Complete; Roche) and phosphatase inhibitors (NaF, Na_4_P_2_O_7_ and Na_3_VO_4_). After preclearing the lysates with Sepharose beads for 15 minutes, antibodies and Protein G Sepharose beads were added and incubated for 2 hours at 4°C. The samples were then washed three times with lysis buffer and once with Tris-NaCl buffer (20 mM Tris pH 7.4, 150 mM NaCl). Samples were boiled in reduced SDS sample buffer and subjected to SDS-PAGE and Western blot as described (Rebeaud *et al.*, 2008).

### Generation of mice expressing non-cleavable Tensin-3

To generate a targeting vector, an approximately 12.7-kb region of the TNS3 gene was first subcloned from a positively identified C57BL/6 fosmid clone (WI1-133G6) using a homologous recombination-based technique. The region was designed such that the 5’ homology arm extends 8.6 kb 5’ to the knock-in mutations. The 3’ homology arm extends 1.4 kb 3’ to the FRT-flanked Neo cassette. The FRT-flanked Neo cassette is inserted downstream of exon 17. The mutations are introduced in exon 17 with the following altering sequences: CGC>GCC for **R**614A and AGA > GCA for **R**645A. The targeting vector was confirmed by restriction analysis after each modification step and by sequencing using primers designed to read from the selection cassette into the 3’ end of the middle arm and the 5’ end of the short arm. The knock-in mutations in exon 17 were confirmed by sequencing. The linearized targeting vector was electroporated into C57BL/6 embryonic stem cells (ES), and positive ES clones were used for injection into C57BL/6 blastocysts (inGenious Targeting Laboratory). To delete the neomycin resistance cassette in the resulting Malt1-KI-neo mice, they were bred to C57BL/6 FLP mice, and Neo deletion was confirmed by PCR. Tensin-3 knock-in and wt mice were genotyped by PCR-mediated amplification of a 391-bp or 277bp fragment, respectively (5’-TGT GGG CCT TTC TGC CTT AAA -3’ and 5’-GTT CCG CCC ACG TCA TAC AC -3’). All mice were maintained on a C57BL/6N background in the specific pathogen-free animal facility of the University of Lausanne. Adult female mice of 8-12 weeks of age were analysed. All mouse experiments were authorized by the Swiss Federal Veterinary Office.

### Mouse immunization

Wt and TNS3 nc/nc littermates (8–12 weeks old) were immunized by intraperitoneal injection with 200 μg of 4-Hydroxy-3-nitrophenylacetyl-chicken gamma globulin (NP-CGG, Biosearch Technology) in alum (Thermo Fisher Scientific). Serum and spleen were harvested for analysis at day 10 after immunization.

### Flow cytometry analysis

Single-cell suspensions from half spleens and inguinal lymph nodes were prepared by gentle mechanical dissociation of respective tissues using a plunger and passage of cells through a 70-μm filter. The bone marrow (BM) was harvested from femurs and tibias, cut at far end and cells were harvested by centrifugation for 1 minute at 13’000 rpm in culture medium (RPMI with 10% FCS). Spleen and BM cells were counted after erythrocyte lysis in ACK buffer (Gibco). Fc receptors were blocked by incubating cells in staining buffer (PBS supplemented with 2% heat-inactivated FCS) with anti-CD16/CD32 antibodies (2.4G2, hybridoma supernatant). Staining was performed in staining buffer on ice for 30 minutes with optimal dilutions. Intracellular staining was performed using the Cytofix/Cytoperm kit (BD Biosciences) according to the manufacturer’s instruction. Viability dye (Invitrogen) was routinely used to exclude dead cells. Data were acquired on a LSR II flow cytometer (both BD Biosciences) and analysed using FlowJo software (TreeStar).

### Isolation of primary B- and T cells

Primary human B and T cells were isolated from healthy donors by MACS beads according to the manufacturer’s description (Miltenyi MACS 130-045-101 for T cells and MACS 130-091-151 for B cells). Isolation of splenocytes and B-cell purification from mice was performed using the B-cell isolation kit according to the manufacturer’s description (Miltenyi MACS 130-090-862).

### In vitro proliferation

2×10^5^ splenocytes labelled with CTV (Invitrogen) were plated in 96-well round-bottom plates (COSTAR) and treated with different stimuli: CpG (3 μg/ml, 1826 Olg ID #833175 from Pascal Schneider), PMA (10 ng/mL; Alexis) and ionomycin (1 μM; Calbiochem) and IgM (10 μg/ml AffiniPure F(ab’)2; Jackson ImmunoResearch). After 4 days CTV dilution was analysed by FACS.

### Adhesion assay

96-well cell culture plates (Sigma) were coated O/N at 4 °C with 50 μl of fibronectin (50 μg/ml; Roche) or VCAM-1-Fc (10 μg/ml R&D System). Coated wells were washed once with PBS (without Ca2^+^ and Mg^2+^) and were blocked for 2 h with 1% BSA in PBS. Cells were left unstimulated or stimulated for indicated time at 37 °C with PMA (10 ng/ml; Alexis) and ionomycin (1 μM; Calbiochem) or solvent (DMSO). Cells were added to the coated tissue culture plates. Subsequently, non-adherent cells were removed by washing of the wells three to five times with with Hank’s balanced-salt solution. Adherent cells were counted using Spectramax (Molecular device) and quantified with ImageJ.

### Detection of serum antibodies by ELISA

For NP-specific antibody detection, Nunc Immuno Plate MaxiSorp plates (Thermo Fisher Scientific) were coated either with 50 μg/ml NP4-BSA or 50 μg/ml NP26-BSA and incubated overnight at 4°C. After blocking with 2% BSA in PBS, the plates were incubated with serially diluted serum samples and then with biotin-conjugated detection antibodies (Southern Biotech) followed by streptavidin-conjugated HRP (Jackson Immunoresearch). After colorimetric reaction (SigmaFAST OPD tablets), the absorbance at 490 nm was measured using an ELISA plate reader. Serum dilutions giving OD50 values for wt sera were chosen. For total Ig detection, Nunc Immuno Plate MaxiSorp plates were coated with 5 μg/ml anti–Ig capture antibody (Southern biotech) and incubated overnight at 4°C. After blocking with 2% BSA in PBS, the plates were incubated with serially diluted serum samples, detected with isotype specific antibodies (Southern Biotech), and developed as described above.

### Xenograft experiments

NSG mice were maintained under pathogen-free conditions in the animal facility of the University of Lausanne. For xenograft studies, NSG mice were inoculated with 10^5^ HBL-1 cells expressing different sgRNAs, mixed with Matrigel (Corning) and PBS in a 1:1 ratio, into the flanks of the animals. Tumor volume was determined by digital caliper measurements and calculated by using the formula (length × width squared)/2. Tumor weight was determined at day 24, when mice were sacrificed.

### Statistical analysis

For statistical analysis prism software was used and two-tailed Student’s *t*-test was applied; *P* values ≤0.05 were considered statistically significant.

## Acknowledgments

The authors would like to thank Benjamin Tschumi and Alena Donda for help with xenograft experiments and initial characterization of the Tensin-3 nc mice, Katrin Bergmann and Anne Müller for advice and initial tests of xenograft experiments, Sergio De Freitas Ribeiro for assistance with the xenograft model, Katherine Clark for Tensin expression constructs, and Fabio Martinon and Marcus Long for discussions and comments on the manuscript. This work was supported by grants to M.T. from the Swiss National Science Foundation (310030_166627), the Swiss Cancer Research foundation (KFS-4095-02-2017-R) and the Emma Muschamp foundation. G.L. was supported by a research grant from the Deutsche Krebshilfe (70113427). S.L. acknowledges support from the Swiss National Science Foundation (310030_185226/1).

## Author contributions

M.J., N.A., I.U. and T.E. designed and performed experiments and analysed data. M.J. and N.A. contributed to writing of the manuscript. M.G. provided technical assistance. G.L. and S.L. designed and analysed experiments and provided feedback on the manuscript. M.T. designed experiments, organized the study, and wrote the paper.

